# The survival of the fastest: Reproductive behavior of the Brazilian three-banded armadillo in the wild

**DOI:** 10.1101/2022.12.12.519664

**Authors:** Nina Attias, Valquíria Cabral Araújo, Carlos Eduardo Fragoso, Amanda de Melo, Guilherme Mourão

## Abstract

The Brazilian three-banded armadillo, *Tolypeutes tricinctus* is endemic to Northeastern Brazil and is classified as a “Vulnerable” (A2cd) species according to the IUCN Red List of Threatened Species. During a study of wild carnivores at Fazenda Trijunção, located at the intersection of the Minas Gerais, Goiás and Bahia states, in Cerrado biome, one of our camera traps recorded three individuals of *T. tricinctus* chasing a fourth one in an area of savanna woodland. This seems to be the record of multiple males competing for access to a receptive female, as a similar behavior is known to occur in some armadillo species. This is the first video record of the chasing behavior for this species, being an indicative that multiple males physically compete to mate with a single female, benefitting fastest males. Information about *in situ* reproductive behavior is essential to contribute to the efficient species conservation planning.

---

Information about *in situ* reproductive behavior is essential to enable efficient species conservation planning (REF). The Brazilian three-banded armadillo, *Tolypeutes tricinctus* (Linnaeus, 1758), is endemic to Northeastern Brazil and is classified as a “Vulnerable” (A2cd) species according to the IUCN Red List of Threatened Species (Miranda *et al*., 2014) and as “Endangered” (A2cd) according to the Official List of Brazilian Threatened Fauna Species (Reis *et al*., 2015). Like many other vertebrates that occur in Northeastern Brazil, the main threats for this species are poaching and habitat degradation (Miranda *et al*., 2014; Reis *et al*., 2015). The creation of a National Action Plan for the conservation of three-banded armadillos (PAN Tatu-bola; ICMBio 2017) further highlights the priority of conservation efforts towards this species. One of the specific goals of the action plan is to broaden the knowledge on the biology and ecology of the species to guide species-specific conservation strategies at a national level (ICMBio, 2017).

We set camera traps to study wild carnivores at Fazenda Trijunção (14°38’34.61”S, 45°48’7.97”W), a private property located at the intersection of the Minas Gerais, Goiás and Bahia states, near the Grande Sertão Veredas National Park, Brazil. This is one of the last refuges for this species’ occurrence in Cerrado biome and likely the southwestern limit of its distribution, being key for this Brazilian endemic armadillo’s conservation efforts (Reis *et al*., 2015). Our camera trap survey started in August 2018 and was composed of 20 camera traps (Bushnell, Trophy Cam HD Brown – 119874, Overland Park, US and Essential E3 Brown-119837, Overland Park, US), set at least 240 meters apart from each other. Cameras were set to record 10-second videos with an interval of 1 second between recordings. The cameras have been monitoring the area continuously, 24h-day, since they were installed (i.e., for the past 19 months), totaling 11,840 camera trap/days.

On June 22^nd^ of 2019 at 2:59 a.m., one of our cameras (14°38’3.87’’S, 45°48’42.73’’W) recorded three individuals of *T. tricinctus* chasing a fourth one (watch video at: https://www.youtube.com/watch?v=Bobnw_n22sg). While one individual ran ahead of the group, the other ones seemed to follow the same approximate path zigzagging and, often, bumping into each other. All individuals were using a trail in an area of savanna woodland. Even though we were not able to capture any of the individuals and we did not record actual intercourse, this seems to be the record of multiple males competing for access to a receptive female, as a similar behavior is known to occur in some armadillo species (Desbiez *et al*., 2006; Tomas *et al*., 2013).

The chasing of a female by multiple males has been previously reported for *T. tricinctus* (Marini Filho & Guimarães, 2010), and for *Euphractus sexcinctus* (Linnaeus, 1758) (Desbiez *et al*., 2006), which was speculatively associated with mating behaviors. Later on, chasing behavior followed by intercourse, was recorded for *E. sexcinctus*, confirming the association of the chasing behavior to the species reproduction (Tomas *et al*., 2013). In addition, males of its congener *T. matacus* (Desmarest, 1804) in captivity have been recorded to chase females before mating events (Meritt, 1976). Hence, most likely this camera trap record should be associated with a reproductive event for *T. tricinctus*.

Our chasing behavior record was made in the month of June, in the same season as the previous report (made in May; Marini Filho & Guimarães, 2010), providing an indication that *T. tricinctus* could present reproductive seasonality in the wild. *T. matacus* presented two peaks of reproduction in the Bolivian Chaco with most pregnant females being detected in the periods of July-September and December-February (Noss, 2003; Cuéllar, 2008). In the Brazilian Pantanal, pregnant females of *T. matacus* have been detected in December-January and July-August, nevertheless, the surveys in this region did not cover every month of the year (Attias N. pers. obs.). The variation on resource availability in the dry forests where *Tolypeutes* occurs may influence the species reproductive period in the wild, although, captive individuals of *T. matacus* have been shown to reproduce year-round (Howell-Stephens *et al*., 2013).

Until now, very little is known about *T. tricinctus* ecology. This is the first video record of the chasing behavior and only the third record of this behavior for the species (Moojen, 1943; Coimbra-Filho, 1972; Marini Filho & Guimarães, 2010). Moreover, the first two mentioned studies have not related the chasing behavior with mating. In contrast with the report provided by Marini Filho & Guimarães (2010), when only two *T. tricinctus* males chased a female, our observation involved three presumed males chasing the presumed female. Actually, Coimbra-Filho (1972) stated that locals reported that this chasing behavior can be performed by even a dozen of armadillos chasing a single one. This chasing behavior is indicative that multiple males physically compete to mate with a single female, benefitting faster males that are capable to detect and keep track of the receptive female, probably through chemical signals. In this case, *Tolypeutes* behavior makes the fastest individuals, the fittest, as biological fitness is given by reproductive success.

Armadillos (Cingulata) present a generally solitary behavior with social interactions restricted to mating and parental care behavior (Ancona & Loughry, 2009; Attias *et al*., 2020). The chasing behavior, associated with *Tolypeutes* space use pattern, where females maintain small exclusive home ranges and males maintain larger home ranges overlapping with both males and females (Guimarães, 1997; Attias *et al*., 2020), provides evidence of a polygynous or promiscuous social system for *Tolypeutes* spp. (Clutton-Brock, 1989), corroborating previous studies for the genus (Attias *et al*., 2020). To further define the species social behavior, we need to understand whether males are capable, or willing, to protect receptive females from other males after the chasing behavior takes place (Attias *et al*., 2020). Hence, behavioral observations will be key to understand this species social system and mating behavior. This information could aid the species conservation *in situ* (e.g., guiding the management of individuals in declining and fragmented populations) and potentially *ex situ* (e.g., promoting husbandry and breeding through the definition of protocols for enclosure sharing during the reproductive season and after cubs are born). Given the steep population decline of *T. tricinctus* in the past decades and the increasing degradation of dry forests and savannas of its restricted distribution range, a “One Plan Approach” with multidisciplinary conservation strategies, integrating both *in situ* and *ex situ* processes (Byers *et al*., 2013), could be adopted and would be benefited by information on the biology of this poorly known species.

## Acknowledgements

We would like thank Pousada Trijunção and Fazenda Trijunção to granting permission to the Onçafari researchers to work on their land. We also thank to Log Nature for providing the Bushnell cameras trap used in this research and the biologist Wellyngton Ayala Espíndola for the fieldwork. We are grateful to the Conselho Nacional de Desenvolvimento Científico e Tecnológico (CNPq) for the fellowship awarded to GM (process 308631/2011-0). This study was financed in part by the Coordenação de Aperfeiçoamento de Pessoal de Nível Superior – Brasil (CAPES) – Finance Code 88887.569720/2020-00.

